# Concurrent ordination: simultaneous unconstrained and constrained latent variable modeling

**DOI:** 10.1101/2021.10.11.463884

**Authors:** Bert van der Veen, Francis K.C. Hui, Knut A. Hovstad, Robert B. O’Hara

## Abstract

1. In community ecology, unconstrained ordination can be used to indirectly explore drivers of community composition, while constrained ordination can be used to directly relate predictors to an ecological community. However, existing constrained ordination methods do not explicitly account for community composition that cannot be explained by the predictors, so that they have the potential to misrepresent community composition if not all predictors are available in the data.
2. We propose and develop a set of new methods for ordination and Joint Species Distribution Modelling (JSDM) as part of the Generalized Linear Latent Variable Model (GLLVM) framework, that incorporate predictors directly into an ordination. This includes a new ordination method that we refer to as concurrent ordination, as it simultaneously constructs unconstrained and constrained latent variables. Both unmeasured residual covariation and predictors are incorporated into the ordination by simultaneously imposing reduced rank structures on the residual covariance matrix and on fixed-effects.
3. We evaluate the method with a simulation study, and show that the proposed developments outperform Canonical Correspondence Analysis (CCA) for Poisson and Bernoulli responses, and perform similar to Redundancy Analysis (RDA) for normally distributed responses, the two most popular methods for constrained ordination in community ecology. Two examples with real data further demonstrate the benefits of concurrent ordination, and the need to account for residual covariation in the analysis of multivariate data.
4. This article contextualizes the role of constrained ordination in the GLLVM and JSDM frameworks, while developing a new ordination method that incorporates the best of unconstrained and constrained ordination, and which overcomes some of the deficiencies of existing classical ordination methods.

## Introduction

Unconstrained ordination methods are used to analyse multivariate data on ecological communities when measurements of the environment are missing. The environment at sites can then only be inferred indirectly from the community composition. For example, when species preferring wet or dry environments are placed at opposite sides of an ordination axis, then this axis will often be interpreted to represent a gradient in soil moisture. This approach of inferring the environment of species relationships can be used to generate new hypotheses (Økland 1996), but by design does not facilitate more exact inference of species relationships due to the environment.

When measures of the environment are available, constrained ordination (also referred to as direct gradient analysis, ter Braak & Prentice 1988) has typically been used in the past to analyse community composition. Constrained ordination is a class of methods akin to multivariate regression, with two notable ones being Canonical Correspondence Analysis (CCA, ter Braak 1986) and Redundancy Analysis (RDA, Rao 1964), which can be used for the analysis of e.g. Bernoulli and Poisson responses, and normally distributed responses, respectively. Constrained ordination arranges sites and species along ordination axes that are constrained to be linear combinations of the measured predictor variables. Species are ordered following the locations of sites, based on their environmental preferences. Constrained ordination uses fewer parameters than (full rank) multivariate regression (ter Braak & Prentice 1988; Yee & Hastie 2003), or its contemporary implementation with stacked (e.g., Wang *et al.* 2012) or Joint Species Distribution Models (JSDMs, Ovaskainen *et al.* 2017), especially when the number of relevant axes is small, as it often is (Halvorsen 2012). For large numbers of predictors and species, a constrained ordination thus leads to a more feasible and interpretable approach for the analysis of multivariate datasets.

In constrained ordination, every added predictor variable provides more flexibility in defining the ordination axis, so that with a large enough number of predictor variables the ecological community as represented by constrained ordination is often quite similar to that represented by unconstrained ordination (Jongman *et al.* 1995; McCune 1997; ter Braak & Šmilauer 2015). An argument for using unconstrained ordination is that the method can be used to explore all variation in the community, whereas constrained ordination filters out variation that is due to the measured environment (Økland 1996) or due to the treatments applied in an ecological experiment (ter Braak & Šmilauer 2015). In situations where few predictors are measured, or when important predictors remain missing, constrained ordination methods such as CCA and RDA misrepresent community composition as any variation not explained by the measured predictors is disregarded (Økland 1996). That is, methods such as CCA and RDA require the assumption that an ordination axis is a perfect function of the predictor variables. However, in practice it can often be unclear which predictors make up an ordination axis, and as such important predictors may remain unmeasured. Consequently, there is a large possibility that there exists covariation between species that cannot be accounted for with the predictors, which invalidates the aforementioned assumption.

The above discussion motivates a unified approach to unconstrained and constrained ordination, which makes it possible to: 1) optimally represent community composition as unconstrained ordination methods do, and 2) concurrently incorporate measured environmental variation with few parameters, as existing constrained ordination approaches do. This requires combining recent advances in model-based unconstrained ordination (e.g., Warton *et al.* 2015) with developments in constrained ordination. Recent developments in model-based unconstrained ordination or JSDMs, as part of the Generalized Linear Latent Variable Model (GLLVM, Skrondal & Rabe-Hesketh 2004) framework are due to Hui *et al.* (2015) for the linear response model, and van der Veen *et al.* (2021) for the quadratic response model, while other authors have made important advances in the form of computational and conceptual developments (e.g., Hui 2016; Ovaskainen *et al.* 2017; Tikhonov *et al.* 2017; Niku *et al.* 2019b; Damgaard *et al.* 2020; Zeng *et al.* 2021). Key developments in statistical models for constrained ordination in recent years include Yee & Hastie (2003) for the linear response model, Yee (2004) for the quadratic (or “Gaussian”) response model (but see also Zhang & Thas 2016), and Yee (2006) for semi-parametric response curves (see also Zhu *et al.* 2005; Hawinkel *et al.* 2019). So far, it has not been possible incorporate residual covariation between species in constrained ordination methods, or to include other kinds of random-effects, but this is exactly the opportunity that the GLLVM framework offers. By combining model-based unconstrained with model-based constrained ordination, it becomes possible to develop new multispecies models with even fewer parameters that utilize the best properties of both methodologies.

We propose a set of new methods for model-based ordination with predictors, that can be regarded as a unifying framework for ordination. Specifically, we propose to relate both predictors and residual covariation to the same latent variable (which we consider to be synonymous to an ordination axis) using a shared set of species-specific parameters. In the new framework, latent variables can be understood as ecological gradients that consist of both measured and unmeasured components. We call the new ordination method developed here *concurrent ordination,* since it simultaneously includes unconstrained and constrained latent variables. Unlike in constrained ordination, the concurrent ordination axes are not necessarily a perfect linear combination of predictors as they can include unmeasured components. This allows the method to discover how well the measured environmental variables are able to model the major structure in community composition.

Through a series of simulations based on normal, Bernoulli, and Poisson species responses, we compare our proposed concurrent ordination approach to two popular constrained ordination methods, CCA and RDA, simultaneously assessing their capability to retrieve the ecological gradients and species responses in the presence and absence of residual variation. We show that in the presence and absence of residual variation, concurrent ordination performs better than CCA in retrieving the ecological gradients and species responses, while performing similarly to RDA for normally distributed responses. Additionally, we demonstrate concurrent ordination with two real datasets; one of alpine plants on an elevation gradient in Switzerland (D’Amen *et al.* 2017), and a dataset of field layer vegetation on an island in Sweden (Cramer & Hytteborn 1987). An easy-to-use software implementation for model-based ordination, with a vignette for more extensive demonstration of the methods proposed here, is available on CRAN as part of the `gllvm R`-package.

## Model-formulation

For a multivariate dataset *y_ij_* consisting of observations recorded for species *j* = 1…*m* and sites *i* = 1…*n*, let *g*(·) generically denote a link function which connects the mean of the assumed response distribution (e.g., the Bernoulli distribution for presence-absence data or the negative binomial for overdispersed counts) to a linear predictor *η_ij_*. We then generally define a GLLVM with linear response model using a vector of ***z***_*i*_ of scores for *d* latent variables and site *i* as:

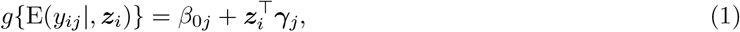

where *β*_0*j*_ is an intercept for each species *j*, and ***γ***_*j*_ is a *d*-dimensional vector of species loadings. For an unconstrained ordination, the site scores are fully formed by residual variation, so that ***z***_i_ = ***ϵ***_*i*_, where we use ***ϵ***_*i*_ to represent a latent-variable level error (“LV-level error” hereafter, in contrast to e.g. the observation-level error usually included in a regression).

For constrained ordination, we also have a vector ***x***_*lv,i*_ of *p* measured predictor variables e.g., solar radiation or available cover, which can also include non-linear terms, that are used to restrict the latent variables in order to filter variation only due to the environment. For a constrained ordination, the site scores are fully determined by predictors, so that ***z***_*i*_ = ***B***^⊤^***x***_*lv,i*_, where ***B*** is a *p* × *d* matrix of slopes that relates the predictors to the latent variables (hereafter referred to as the “canonical coefficients”). With categorical predictors, constrained ordination clusters sites, so that the canonical coefficients are measures of distance in the ordination space between the groups. For continuous predictors, constrained ordination orders sites instead.

Compared to standard multivariate regression, constrained ordination with a linear response model reduces the number of parameters, for which the parameters can be especially difficult to estimate with large *p* and small *n* (i.e., with many predictors and a small number of sites). The vector of coefficients per species *j* from a multivariate regression *β_j_* can be reconstructed from a constrained ordination as ***β***_*j*_ = ***Bγ***_*j*_ with accompanying standard errors (see Appendix S1). For *d* = min(*m, p*) the constrained ordination includes as many parameters as a multivariate regression, namely *m* + *mp*. However, when *d* < *p* ≪ *m*, constrained ordination includes fewer parameters than a multivariate regression, which can often be a more practically appropriate assumption for ecological community data that tend to be sparse on information. Specifically, the number of parameters in constrained ordination is *m* + *d*(*m* + *p*) – *d*^2^ (Robinson 1973) for a rank *d* solution i.e., with *d* latent variables.

The formulation of constrained ordination outlined above shows that it disregards any variation that can not be explained by the predictors. To overcome this, concurrent ordination instead constructs latent variables that are still assumed to be a function of the predictors, but also includes the LV-level error term 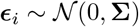 from unconstrained ordination, where we assume **Σ** = diag(***σ***^2^), i.e. independent LV-level errors for sites and latent variables. Formally, the proposed model for concurrently performing unconstrained and constrained ordination is:

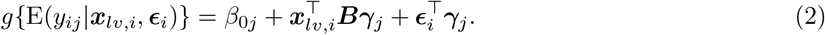

The model in equation (2) can also be formulated in terms of the latent variables ***z***_*i*_ = ***B***^⊤^***x***_*lv,i*_ + ***ϵ***_*i*_, including two reduced-rank terms that are linked together by the vector of species loadings ***γ***_*j*_. As such, concurrent ordination includes constrained ordination as a special case with var(***ϵ***_*i*_) = **0**, and unconstrained ordination when ***B*** =**0**. In situations that the predictors do not fully explain the latent variables, and there is residual variation present, concurrent ordination both describes the effects of important predictor variables in shaping community composition, while concurrently simplifying the interpretation (“reify”) of the ordination.

By noting that in concurrent ordination we impose a rank constraint for the the matrix of predictor slopes, in addition to a rank constraint on the residual covariance matrix, and that the imposed rank of those two matrices is the same, we can alternatively interpret the model in equation (2) as either: 1) having connected sets of constrained and residual latent variables, or 2) as a special case of model-based residual ordination as in typical JSDMs (see e.g., Hui *et al.* 2015; van der Veen *et al.* 2021). Following the developments in van der Veen *et al.* (2021), the ordination methods presented here can also be extended to the quadratic response model:

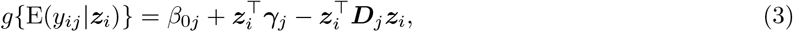

where ***D***_*j*_ is a positive-definite diagonal matrix that contains the quadratic coefficients for the latent variables per species. Setting again ***z***_*i*_ = ***ϵ***_*i*_ for an unconstrained ordination as in van der Veen *et al.* (2021), ***z***_*i*_ = ***B***^⊤^***x***_*i*_ for a constrained ordination as in Yee (2004), and ***z***_*i*_ = ***B***^⊤^ ***x***_*i*_ + ***ϵ***_*i*_ for a concurrent ordination. For the latter, writing out the model completely will reveal its hidden and high degree of complexity, so that the assumption of species-common tolerances, i.e., ***D***_*j*_ = ***D*** is often more realistic. More detailed discussion of this extension for concurrent ordination to the quadratic response model is provided in Appendix S2.

All of the aforementioned models can be extended to include additional predictors ***x***_*i*_, separate from those in the ordination ***x***_*lv,i*_, resulting in e.g. a partial concurrent ordination, similar to ter Braak (1988) and reduced rank regression with concomitant variables (Davies & Tso 1982):

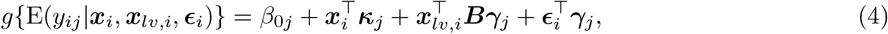

where ***κ***_*j*_ are species coefficients for the predictors ***x***_*i*_, and where we additionally assume that ***x***_*i*_ and ***x***_*lv,i*_ do not include the same predictor variables for reasons of parameter identifiability. Here, the effect of ***x***_*i*_ is excluded from the concurrent ordination, so that the resulting ordination is interpreted conditionally on the predictors ***x***_*i*_ (referred to as “conditioning” in classical ordination methods, see e.g. Hawinkel *et al.* 2019). All of the models are supported by the `gllvm R`-package for implementation (Niku *et al.* 2020), and can be straightforwardly extended with additional random intercepts to e.g. account for pseudoreplication of sites, or to model community composition instead, though we have chosen to omit that term here for ease of presentation.

## Parameter identifiability

Constrained and concurrent ordination are unidentifiable without parameter constraints. To impose constraints, consider a *p* × *d* matrix **Γ** that includes all species loadings ***γ***_*j*_ as row vectors, for which we fix all entries above the main diagonal to zero, as is usual for GLLVMs (Hui *et al.* 2015).

In the standard formulation of GLLVMs, to account for scale invariance, the latent variables are assumed to have unit variance. Then, the columns of the matrix species loadings **Γ** regulate the scale of the ordination. For concurrent ordination as formulated in equation (2), the species loadings are shared for two terms, so that without separating the scale for the latent variables from the species loadings, the columns of ***B*** will regulate the relative scale of the second and third term in (2). As an example, in cases where the fixed-effects term is non-zero and there should be no LV-level error (i.e., when it is zero), but a concurrent ordination is fitted regardless, the loadings are required to be very small as they simultaneously regulate the scale of both terms. This can then only be compensated for by increasing the magnitude of the canonical coefficients ***B***, potentially resulting in numerical issues. Therefore, we additionally choose to fix one parameter in the species loadings per latent variable to facilitate including freely varying scale parameters ***σ***^2^ for the LV-level error ***ϵ***_*i*_. Here, we choose to fix the parameters on the main diagonal of **Γ** to one, such that in **Γ** there are only *md* – *d*(*d* + 1)/2 parameters to estimate. This choice of the diagonal elements is arbitrary, and different elements could be chosen instead. This current choice is guided by the reasoning that now ***B*** always determines the scale of the first (fixed-effects) term, so that it is (close to) zero when the predictors have no effect on the ordination, and non-zero otherwise. Similarly, the vector of standard deviations ***σ*** then reflects the scale of the LV-level error, so that it is zero when there is no LV-level error necessary in the ordination (i.e. when the latent variables are perfectly represented by the predictors, as assumed in constrained ordination), and so that the model can determine the relative contribution of the measured and unmeasured components based on the data. Note that this choice of identifiability constraint does not diminish the overall flexibility of the model, but merely clarifies the interpretation of the parameters: in essence, the latent variables are stretched or contracted so that the species loadings on the main diagonal equal one. So far, the number of identifiability constraints is the same as in a model-based unconstrained ordination (Hui *et al.* 2015), and additional parameter constraints are required to fully identify the proposed constrained and concurrent ordination models. Yee & Hastie (2003) further fixed additional parameters in **Γ**, but we choose a different parameterization. We instead require the columns of ***B*** to be orthogonal, i.e. for a diagonal matrix ***C*** with entries equal to the columnwise Frobenius norms of ***B***, we require ***B***^⊤^ ***B*** = ***C***^⊤^***C***, thus adding *d*(*d* – 1)/2 parameter constraints. We note that following this parameterization, for a linear response model, and in particular when *d* = *p*, ***B***Γ^⊤^ is the estimated QR-decomposition of the matrix of predictor slopes in a multivariate regression, with ***Q*** = ***BC***^-1^ and ***C***Γ^⊤^, and its estimated reduced QR-decomposition when *d* < *p*. Finally, we note that as in the VGAM `R`-package, the model can be sensitive to the order of species in the data occasionally, so that if convergence issues arise this can in some cases be improved by re-ordering the columns of the response data, or equivalently by rotating the ordination.

## Parameter estimation

For the proposed methods we are required to choose an appropriate distribution, with associated mean-variance relationship, to model the species observations. For example, a Poisson or negative-binomial distribution with log-link function may be used for count data, a Bernoulli distribution with probit link-function may be used for binary data, or alternatively a Tweedie distribution with log-link function may be used for biomass data. Because the LV-level error ***ϵ***_*i*_ are assumed to be random variables, they needs to be integrated over. Consequently, the marginal log-likelihood of the proposed concurrent ordination methods, as in equation (2), is written as:

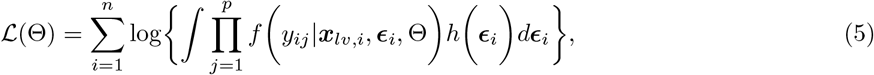

where *f* (*y_ij_*|***x***_*lv,i*_, ***ϵ***_*i*_, Θ) is the distribution of the responses conditional on the predictors ***x***_*lv,i*_, the LV-level error ***ϵ***_*i*_, and a vector of parameters Θ that includes all freely varying model parameters. The LV-level error is assumed to follow a multivariate normal distribution, 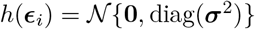. The integration can be performed with the approaches available in the `gllvm R`-package previously developed for approximate estimation and inference in GLLVMs for unconstrained ordination (Niku *et al.* 2019a), namely the Laplace approximation (Niku *et al.* 2017) or Variational Approximations (VA, Hui *et al.* 2017; van der Veen *et al.* 2021), and implemented via Template Model Builder (Kristensen *et al.* 2016). The LV-level residual can be obtained as e.g., the means of variational distributions (Hui *et al.* 2017) or the maximum a-posteriori prediction from the Laplace approximation (Niku *et al.* 2017). Below, the models in the simulation studies and examples are fitted using VA.

The orthogonality constraints on the canonical coefficients require maximizing equation (5) subject to the equality constraint ***B***^⊤^***B*** – ***C*** = 0. To do this, we make use of the augmented Lagranian method implemented with the `alabama R`-package (Varadhan 2022), which allows maximization with (in)equality constraints using `R`’s native optimization routines `optim` and `nlminb`. In practice we found that this generally worked out well, so that it was usually possible find a local maximum of the objective function, but since there are situations in which the use of alternative optimization routines can aid in diagnosing and addressing convergence issues (Bates *et al.* 2014), we additionally provide support for two alternative optimization routines. Specifically, we provide access to another implementation of the augmented Lagrangian method, and to a sequential quadratic programming method, both from the NLopt library of non-linear optimizers (Johnson 2014) and implemented via the `nloptr R`-package (Ypma *et al.* 2018).

### Initial values

Both with and without LV-level error, the algorithm used to fit the models presented here is sensitive to the initial values. In this article, we adapt the approach used in the `gllvm R`-package to overcome this, and obtain reasonable starting values. Specifically, for constrained ordination, we followed a similar procedure to that described by Files *et al.* (2019), where we generate starting values for ***B*** and ***γ*** by first fitting a multivariate linear model with predictor variables to the Dunn-Smyth residuals (Dunn & Smyth 1996) of an intercept-only multivariate Generalized Linear Model. We then performed a QR-decomposition on the matrix of regression coefficients, from which we take the first *d* dimensions, to obtain the starting values for ***B*** and ***γ***_*j*_. For concurrent ordination, and when *m* < *n* we perform a maximum likelihood factor analysis on the Dunn-Smyth residuals of an intercept-only MGLM, and then regress the estimated factor scores to receive initial values for the canonical coefficients ***B***. Otherwise, we transpose the data and follow the same procedure, but regress the loadings instead. The residuals of the regression can then be used to initiate the algorithm, i.e. as the starting values for the estimate of E(***ϵ***_*i*_|***y***_*i*_), and the loadings from the factor analysis can be taken as the initial values for ***γ***_*j*_.

## Inference

In this section, we present various tools for inference and prediction for the proposed concurrent ordination method.

### Ordination diagram

Generally, an ordination diagram for model-based ordination can be interpreted similarly as that for a classical ordination. In the case of concurrent ordination, ordination diagrams can be constructed for the second and third terms in equation (2) to explore species co-occurrence patterns due to the predictors and residual covariation separately, or a single ordination diagram can be constructed including both terms. Note that since the latter ordination includes LV-level residuals, it is important to assess the relative contribution of predictors to the latent variables (see also the section on *Predictor importance* below) since site and species coordinates are not required to (only) exhibit patterns that relate to the predictors.

Unconstrained latent variables are assumed to fully consist of residual information, as discussed previously. By contrast, in concurrent ordination, the predicted site scores ***z***_*i*_ can be constructed based on the predicted ***ϵ***_*i*_’s along with the ***B***’s. We define separate sets of: 1) conditional scores ***B***^⊤^***x***_*i*_ + ***ϵ***_*i*_ that compare to weighted average or weighted summation scores in CCA and RDA, respectively 2) marginal scores ***B***^⊤^***x***_*i*_ that do not include an estimate of the LV-level error, which correspond to linear combination scores for classical constrained ordination methods, and 3) residual scores ***ϵ***_*i*_ that are unique to the methodology presented here and do not include effects of the predictors. The scores can be used to check the fit of the predictors to the latent variables, e.g. to check linearity (plot of conditional scores versus specific predictors) or heteroscedasticity assumptions (plot of residual scores versus marginal scores), similar as in residual diagnostics for ordinary linear regression.

Since marginal scores do not include additional information on the latent variable provided by the response data, they are not generally recommended for inference in community ecology (McCune 1997). We consider conditional site scores similar to the weighted average or weighted summation scores (Palmer 1993; McCune 1997) from CCA or RDA, because those scores can be considered as minimally constrained, and since the LV-level error from our model ***ϵ***_*i*_ accounts for variation in the response not explained by the predictor variables. To summarize, a separate ordination diagram can be drawn for conditional site scores, marginal site scores, or residual site scores, depending on the effects a researcher wishes to emphasize. For new measurement data of the environmental variables, and if no observations of species at the new sites are available, one can calculate the associated marginal scores, but the conditional scores are not available.

An ordination diagram with conditional site scores will, in many instances, provide a similar ordination as when latent variables are assumed to be unconstrained. However, the predictor effects can now be represented in the ordination diagram as in constrained ordination, in the form of arrows based on the rows of ***B***. The length of each arrow is proportional to the magnitude of the parameter estimate, so that the predictor with the largest estimate is presented as the longest arrow, although note that we correct the arrow length using the standard deviation of each predictor (see Figure 3 below). Statistical uncertainty of the estimated canonical coefficients for the predictors can be further represented using the colour of the arrows, for example by colouring the arrow less intensely for estimates with a confidence interval that includes zero for at least one of the plotted ordination dimensions.

In an ordination diagram, the predicted site scores (irrespective of which of the three versions of site scores introduced above) are plotted to represent (dis)similarity between sites in an ordination. Furthermore, Niku (2020) constructed corresponding prediction regions using the Conditional Mean Squared Error of Predictions (CMSEPs, Booth & Hobert 1998) to represent the statistical uncertainty of the site scores in an ordination diagram. To fully and properly convey confidence in the dissimilarity of sites, we adopt the same approach, but modify the calculation for the case of concurrent ordination (see Appendix S3 for details of the calculation). These prediction intervals can be used to provide a larger degree of certainty in the position of sites in an ordination, and include both the statistical uncertainty of the fixed-effects and of the prediction for the LV-level error.

### Model and variable selection

As the number of measured predictors increases, and more variation in the latent variable is accounted for by the predictors, the standard deviations of the LV-level error ***σ*** are likely to get smaller. Determining the optimal number of latent variables and the most suitable predictor variables for an ordination is thus an important problem for concurrent ordination, and it can be a challenging exercise as the number of potential models may be quite large. Fortunately, in a model-based framework as proposed here, it is possible to leverage conventional methods such as hypothesis testing, information criteria (Burnham & Anderson 2002), and residual diagnostics among others for assessing the optimal number of predictors, for predictions, as well as for assessing other model assumptions such as the distribution of the responses. For example, the importance of predictors in a concurrent ordination can be assessed with use of a partial *R*^2^-statistic. We illustrate an example of determining predictor importance later on in our applications of two datasets of real ecological communities. The number of latent variables supported by the data, can be solved using model selection tools such as information criteria (Skrondal & Rabe-Hesketh 2004 p. 266; Bartholomew 2011 p. 58), or using regularization techniques (Hui *et al.* 2018).

As an alternative to the above, it is possible to utilize a random-effects formulation in order to penalize the canonical coefficients while also automatically selecting the corresponding tuning parameter (Robinson 1991). Predictor effects in constrained ordination methods are often unstable (i.e. have increased variance, ter Braak 1994). Treating the canonical coefficients ***B*** as random-effects and shrinking them will reduce their variance and has the potential to stabilize estimation (see Appendix S4). Since for categorical predictors the canonical coefficients are intercepts, while they are slopes for continuous predictors, this development implies to the inclusion of both random intercept and random slope terms in a constrained or concurrent ordination. The covariance structure of these random-effects can be assumed to be latent variable specific in order to additionally induce correlation between the responses of species to the same predictor, or predictor specific in order to shrink some predictor effects to near zero (see Appendix S4 for further discussion on this). The coefficients of predictors that are less important are shrunk by a larger degree, thus reducing their role in the model. Although random-effect shrinkage cannot shrink coefficients to exactly zero, in practice the coefficients of predictors that are not supported by the data will be sufficiently close to zero, so that this method can be used for variable selection (e.g., by adopting a sort of thresholding). Methods for random-effects shrinkage are supported in the the `gllvm R`-package for both constrained and concurrent ordination. Note that in the simulations and examples below, we assume that the canonical coefficients are fixed-effects.

### Predictor importance

Similarly to other GLLVMs, the residual covariance matrix associated with the latent variables can be calculated (see Appendix S5 for details). This residual covariance matrix allows researchers to examine species associations as in other JSDMs, and to determine the residual covariation in the response beyond that due to the measured predictors. Furthermore, by first fitting a concurrent ordination to the data, and secondly an unconstrained ordination, the variation explained by the predictors in the response can be determined based on relative differences in the trace of the residual covariance matrices of both models (Warton *et al.* 2015).

Since the variation explained by predictors in the response is not necessarily a good measure of their importance in an ordination, we here focus on determining the importance of predictors in explaining the latent variables instead. Note that the latent variables are determined by a linear regression, nested in the multivariate model. Since the latent variables are by definition (partially) unmeasured, calculating importance of the predictors through e.g. a partial *R*^2^, as in ordinary linear regression, is not possible. As such, to assess the importance of predictors in explaining the latent variables, we adopt an approach similar to that presented by Edwards *et al.* (2008), which avoids having to fit a second model (see Appendix S6). Specifically, Edwards *et al.* (2008) developed a measure of *R*^2^ for linear mixed-effects models based on the fit of a single model, which Jaeger *et al.* (2017) extended to the generalized linear mixed-effects model and implemented in the `r2glmm R`-package, using a multivariate Wald-statistic for the testing of fixed-effects.

We will refer to this specific (semi-partial) *R*^2^ here as 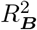. To be clear, 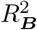 measures the importance of the predictors in explaining the latent variables, not the importance of the predictors in explaining the response data (though that could similarly be calculated). Note that 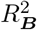 can also be calculated on a per predictor variable basis (with numerator degrees of freedom *d*), or per latent variable and predictor (with unit numerator degrees of freedom). The correct interpretation for 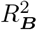 is the (generalized) residual variation a particular predictor can account for, after accounting for all other predictors in the model (Edwards *et al.* 2008). We demonstrate use of 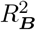 in the real data examples below. Note that calculating 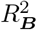 for a fitted constrained ordination has the potential to generate artificially high values if there is in reality residual variation unaccounted for, and in particular when little statistical uncertainty is associated to the parameter estimates.

To assess importance of predictors when the canonical coefficients are treated as random-effects instead, the scale parameters for the canonical coefficients and for the LV-level error can be used to construct a *R*^2^ similar to e.g. Nakagawa & Schielzeth (2013).

## Simulation studies

To assess empirical performance of the proposed GLLVM, we simulated from models with and without LV-level error (i.e., concurrent and constrained ordinations), and examined the capacity to retrieve the true latent variables ***z***_*i*_ and species loadings ***γ***_*j*_. We then compared performance to CCA and RDA, applied with the `vegan R`-package. We provide `R`-code for reproducing all the simulations in Appendix S7.

To be more precise, we generated data based on two forms of the model in equation (2): with 1) non-zero canonical coefficients ***B*** and LV-level error ***ϵ***_*i*_ i.e. a concurrent ordination, 2) non-zero canonical coefficients while ***ϵ***_*i*_ = 0, i.e. a constrained ordination. We considered datasets with *n* = 100 sites and with *m* = 30 species, since it is more difficult to accurately predict the LV-level error when the number of species is small.

To construct the true model, we first simulated *p* = 5 predictor variables following a multivariate standard normal distribution. Next, we generated the true canonical coefficients ***B*** as the loadings from a maximum likelihood factor analysis fitted to the simulated predictor variables with two dimensions. We simulated the true intercepts *β*_0*j*_ independently from Uniform(–1,1), and species loadings ***γ*** independently from Uniform(–2, 2). Finally, we simulated the LV-level error ***ϵ***_*i*_ by sampling from a bivariate standard normal distribution, after which we regressed the sampled realization against the simulated predictor variables, and used the (to unit variance scaled) residuals from the regression as the LV-level error in the true model. This ensures that the true LV-level error ***ϵ***_*i*_ was independent of the simulated predictor variables by construction. The two forms of true model were then formed based on whether ***ϵ***_*i*_ was omitted from the model or not. For each of the two possible models, and for Normal, Bernoulli, and Poisson species responses, we simulated 1000 datasets. The variance associated with Gaussian responses was assumed to be one. To each dataset, we fitted a concurrent ordination with two latent variables. As a comparison, we applied CCA to all simulated datasets with Bernoulli and Poisson species responses and applied RDA all simulated datasets with Gaussian distributed responses. To assess performance, we calculated the Procrustes error between the simulated latent variables and the latent variables retrieved from the proposed GLLVM, CCA, and RDA, and the same for the species loadings (Peres-Neto & Jackson 2001; Oksanen *et al.* 2020). When constrained ordination was the true model, we used linear combination scores for CCA and RDA, and when concurrent ordination was the true model, we used weighted average or summation scores. The results of the simulations are summarized in Figure 1.

**Figure 1:**
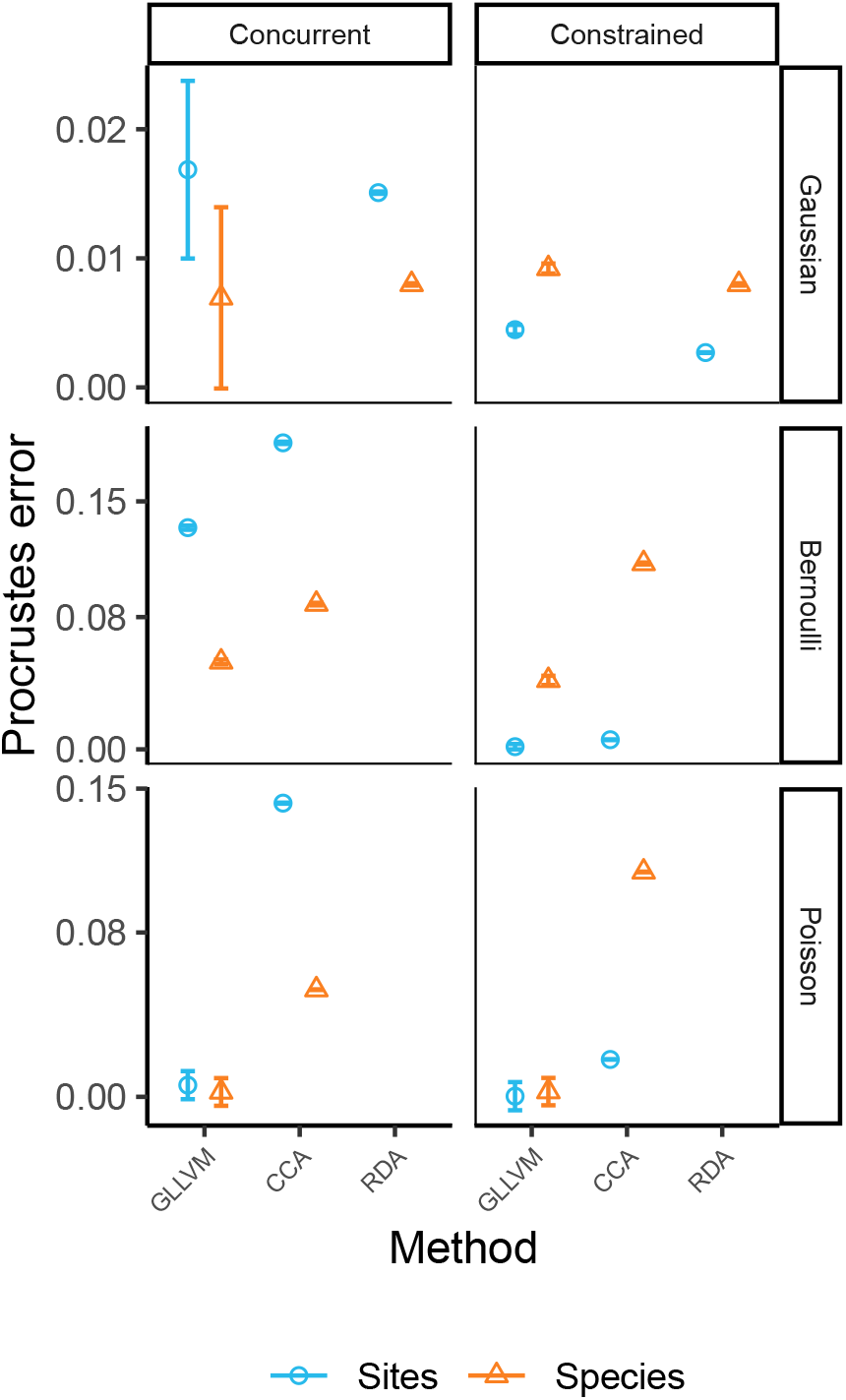
Results for ordination methods fitted to 1000 simulated datasets with normally distributed, Bernoulli, and Poisson species responses. For simulations with columns labelled ‘constrained’, constrained ordination was the true model (i.e. we assumed the LV-level error was fixed to zero ***ϵ***_*i*_ = **0**), and for rows labelled ‘concurrent’, concurrent ordination was the true model. The mode of the Procrustes error for the latent variables ***z***_*i*_ is shown in lightblue and with a circle, and for the species loadings ***γ*** in orange and with a triangle, with error bars representing the 95% confidence interval.

**Figure 2:**
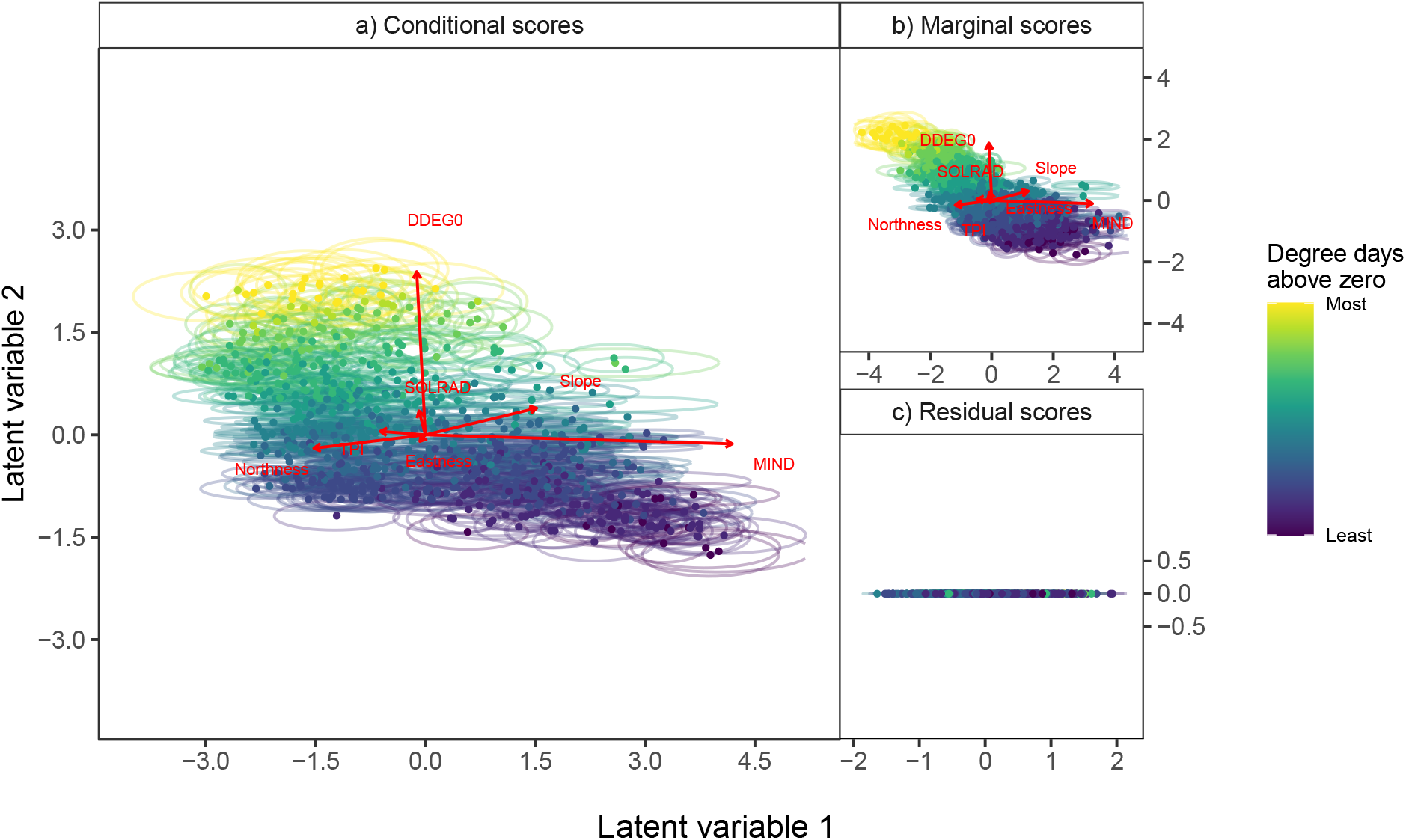
Ordination diagrams for the Swiss alpine plants data. Darker colors indicate sites with fewer degree days above zero, whereas lighter colors indicate sites with more degree days above zero. Arrows represent predictor effects for each latent variable, with arrow length being proportional to the magnitude of the canonical coefficients estimates for each of the predictors: Degree days above zero (DDGEG), eastness and northness, moisture index (MIND), slope, total solar radiation over the year (SOLRAD), and topograhy index (TPI). Each plot includes a separate set of site scores; conditional (a), marginal (b), and residual scores (c) respectively. Ellipses represent 95% prediction regions. No arrows are drawn for the residual scores, since the LV-level error is by design uncorrelated with predictor effects, and arrows would serve no purpose. Detailed results, and a list of names for species included in the data, are included in Table S1 and Table S3 of Appendix S8.

In general, GLLVMs managed to retrieve the true latent variables ***z***_*i*_ and species loadings ***γ***_*j*_ consistently and with little variability. Similarly, for Gaussian responses, RDA managed to retrieve the latent variables and species loadings equally well. In all cases, GLLVMs performed better than CCA.

## Worked examples

We demonstrate applications for the proposed concurrent ordination method on two ecological datasets: 1) a dataset of Swiss alpine plants (D’Amen *et al.* 2018), and 2) a dataset of field layer vegetation on the island Skabbholmen in Sweden (Cramer & Hytteborn 1987).

### Swiss alpine plants

The first example focuses on a presence-absence dataset of alpine plants, collected in the western Swiss Alps. The dataset was collected on a strong elevation gradient, including sites in both lowland and alpine environments (D’Amen *et al.* 2018). In total the dataset includes *m* = 175 plant species and *n* = 791 plots, after excluding plots with fewer than two species, and species with fewer than three presences. Seven predictor variables were included in the study: degree days above zero, slope, moisture index, total solar radiation over the year, topography, and aspect (as northness and eastness). All predictors were scaled to have unit variance, and centered to have mean zero, prior to fitting. We additionally applied CCA to the data and calculated the Procrustes error between the marginal and conditional site scores from the fitted concurrent ordination, and the linear combination and weighted average scores from the fitted CCA, to determine how similar the solutions of the two methods were.

Fitting a range of models while testing for the optimal number of dimensions and predictors would be computationally burdensome and time consuming. Therefore, we focused on fitting a single model with *d* = 2 latent variables, using all predictors, and with one quadratic coefficient per latent variable (i.e. common tolerances) for demonstration purposes. Previously, van der Veen *et al.* (2021) found that using a quadratic response model lead to better predictions of the ecological gradient for this dataset, so we adopt the same approach here. Additionally, since the data are binary, we fitted the model with a Bernoulli distribution. We then base our inference on the confidence intervals of the canonical coefficients, Wald-statistics with accompanying P-values, and the approach for 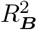 presented above.

The results are visually presented in Figure 3. More detailed information from the ordination is available in Table S1 of Appendix S8. Similar to van der Veen *et al.* (2021), degree days above zero was the predictor most related to the two predicted latent variables.

**Figure 3:**
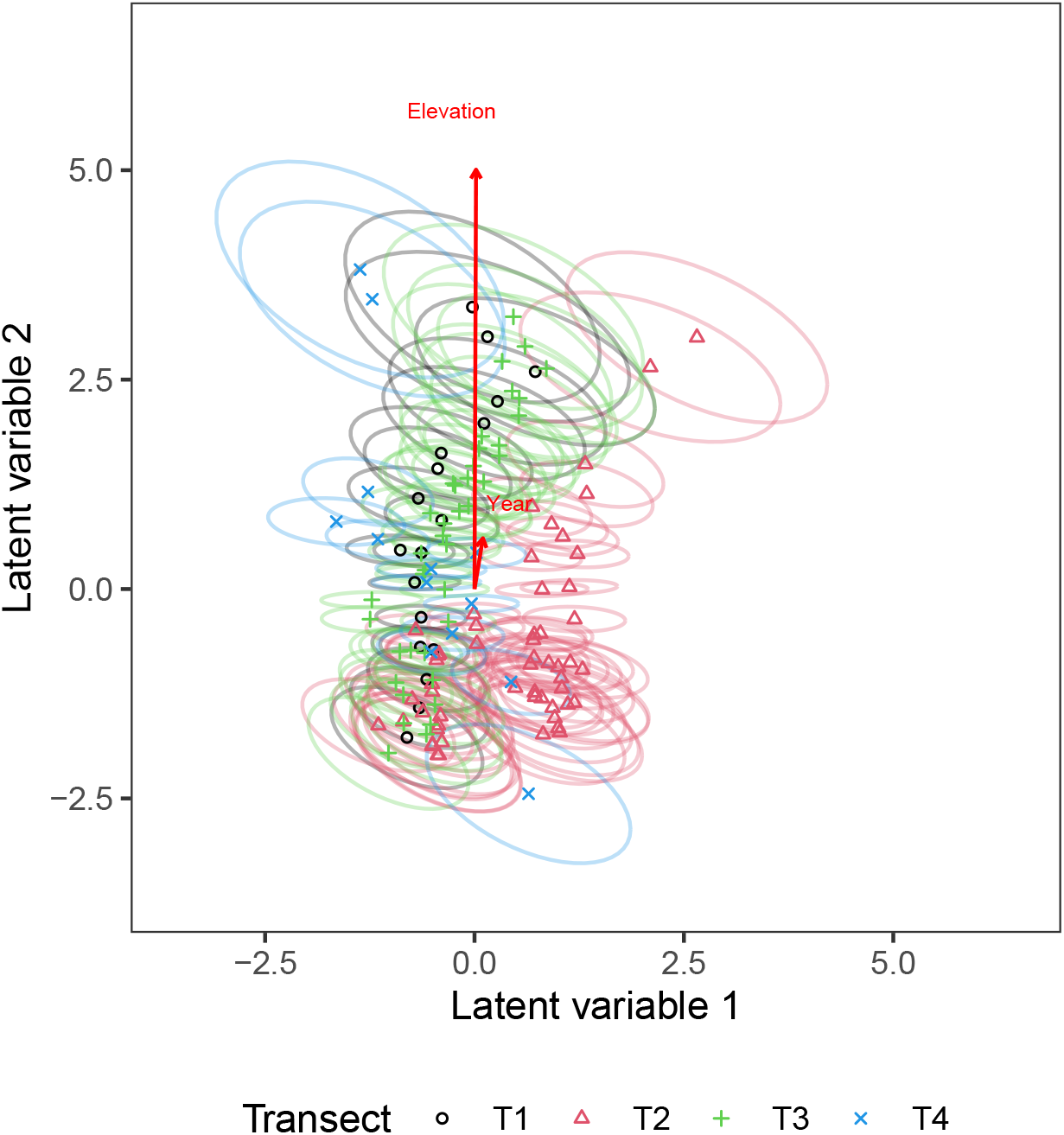
Concurrent ordination plot for sites from the Skabbholmen dataset, conditional on a random row intercept to account for pseudoreplication of transects. Sites with corresponding 95% prediction regions are colored by their respective transects. Estimates and standard errors of the canonical coefficients are provided in Table S4 of Appendix S8.

Based on the fitted GLLVM with two latent variables and containing all six predictor variables, the estimated standard deviations of the LV-level error were 2.10 (95% confidence interval: 1.78, 2.43) and 0.00 (0.00, 0.01), indicating that the second latent variable could be fully represented by the predictors. Mixed-effects models with scale parameters on the boundary of the feasible parameter space usually suffer from numerical issues (Oberpriller *et al.* 2022), and in a realistic workflow the model should have been refitted while omitting the LV-level error for the second latent variable, but we continue with the same model here for illustrative purposes. The zero variance associated with the LV-level error of the second latent variable is also visually indicated by Figure 3c, where the points lay approximately parallel to the x-axis as a consequence. The 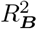 for the latent variables was 0.17, while for the predictors this was 0.10 (degree days above zero), 0.09 (slope), 0.04 (solar radiation), 0.03 (northness), 0.02 (eastness), 0.02 (moisture index), and 0.01 (topography), indicating that degree days above zero and slope were the most important predictors for representing the latent variables. Similar for the concurrent ordination, CCA had a low 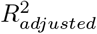 of 0.11 (see also Figure S1 and Table S2 in Appendix S8). The Procrustes error, with both solutions scaled to equal dispersion, between the marginal site scores of the concurrent ordination and the linear combination scores from CCA was 0.25. The Procrustes error between the conditional site scores of the concurrent ordination and the weighted average scores from CCA was 0.43.

### Field layer vegetation on Skabbholmen, Sweden

The second example includes ordinal responses on a five-degree Hult-Sernander-Du Rietz scale (Du Rietz 1921; Oksanen 1976). Vascular plant cover was recorded in *n* = 135 one-square-meter plots unequally divided over four transects (Cramer & Hytteborn 1987). Cramer & Hytteborn (1987) split the data based on a clustering algorithm, and only the sites with shoreline vegetation are included in the data here, bringing the number of unique sites to *n* = 64, of which two sites were sampled in only one year (see also Jongman *et al.* 1995 p. 167 for more information). We excluded species that occurred in two plots or less, such that the final number of species in the analysis was *m* = 49. Each transect was recorded in two different years (1978 and 1984), and followed an elevation gradient from the shoreline to the edge of old-growth forest, but note that elevation (in centimeters) was only recorded during the first sampling. The elevation gradient serves to represent long term effects of land uplift as a consequence of the retraction of land-ice in Scandinavia after the last glacial maximum, whereas the year effect serves to represent the actual temporal change in community composition. Both predictors were scaled to unit variance and centered to mean zero prior to model fitting.

Originally, the data was analyzed with a combination of Detrended Correspondence Analysis (Hill & Gauch 1980), CCA, and detrended CCA, leading the authors to conclude that elevation was the main driver of community composition. ter Braak (1987) used the ratio of the two canonical coefficients for the dominant axis to determine if the change in vegetation tracks the known land uplift of half a centimeter a year. The resulting ratio coefficient was 0.76, and although this was higher than the known land uplift, the approximate confidence interval included the known land uplift. For comparison, we here calculate the same ratio, and also provide an approximate 95% confidence interval calculated using Fieller’s theorem (see Lye & Hirschberg 2018, equation 10).

We fitted a concurrent ordination with *d* = 2 latent variables to the data, and also included random row intercepts to account for the pseudoreplication of transects (Cramer & Hytteborn 1987 wrote in the footnote on page 164 that residual variation could be attributed to the difference in transects), assuming a multinomial distribution with cumulative probit-link function for the response, and with species-common tolerances (i.e., one quadratic coefficient per latent variable). In this concurrent ordination, the estimated standard deviation for the second LV-level error was near zero and so we re-fitted the model omitting the LV-level error for one latent variable. Consequently, the estimated standard deviation for the LV-level error of the first latent variable was 1.33 (95% confidence interval: 0.69, 1.97), and the canonical coefficients for the first latent variable were near zero (see Table S4 in Appendix S8). The 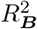 for the latent variables was 0.27, while for the predictors this was 0.27 (elevation) and 0.07 (year), indicating that the year effect was not an important factor in explaining the latent variables. In summary, the first latent variable was unrelated to the predictors but the second latent variable was nearly fully explained by elevation.

The results for this model are visualized in Figure 3, where sites that come from the same transect have been indicated with the same symbol and color. Species optima have been omitted, because species responses for the first latent variable were nearly linear (i.e., ***D*** ≈ 0), but the species optima for the second latent variable are instead visualized in Figure 4. Since elevation almost fully explained the second latent variable, and since the first latent variable was unrelated to the predictors, vertical separation in the plotted site scores can be considered due to elevation, whereas it remains unclear what drives the horizontal separation in the site scores. Consequently, large positive values of the second latent variable correspond to species or sites found near the forest edge, and large negative values to species or sites found at lower elevations, i.e., at the shoreline.

**Figure 4:**
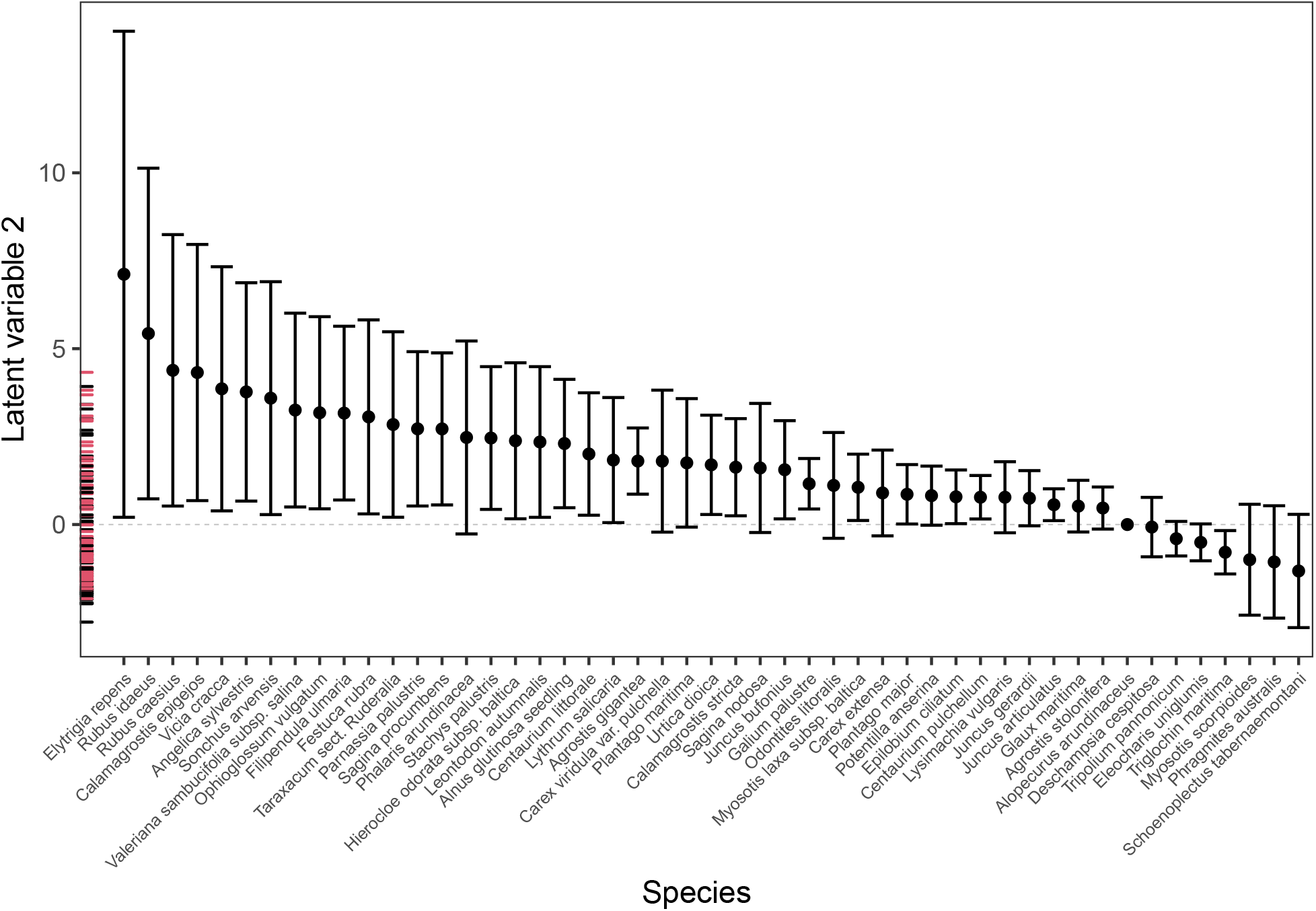
Concurrent ordination plot of species optima for the first latent variable. The latent variable almost completely corresponds to elevation, so that species optima with high values represent species preferring (relatively) higher elevations, and species optima with negative values lower elevations. Large positive values of the latent variable thus correspond to species or sites found closer to the forest edge, and large negative values to species or sites found closer to the shoreline. Error bars represent approximate 95% confidence intervals. A dashed line has been added as a visual aid at zero, to which the optimum of the first species on the second latent variable is fixed for reasons of parameter identifiability (fixing the rotation of the ordination). Site scores are indicated by a rugplot on the y-axis, colored by sampling year, 1978 corresponding to black and 1984 to red.

Finally, the ratio of the year and elevation coefficients was 0.77 (approximate 95% confidence interval based on a t-distribution with *n* – 2 degrees of freedom: 0.52, 1.01). This ratio of parameter estimates, and its confidence interval, are nearly the same as reported by ter Braak (1987), but here the confidence interval excludes the known land uplift of 0.5 centimeter a year, thus leading to a different conclusion than ter Braak (1987) and Cramer & Hytteborn (1987), namely that vegetation change was faster than the known land uplift.

## Discussion

In this article, we present new methods for ordination with predictor variables, or alternatively for estimating species responses in a reduced-rank form, with GLLVMs. Unconstrained ordination allows ecologists to order sites and species when the environment is unmeasured, while constrained ordination restricts the ordering using measured predictors of the environment, in order to better examine species-environment relationships (ter Braak 1987). The framework for ordination proposed here goes beyond these concepts by combining the properties of both unconstrained and constrained ordination, and is rooted in the social sciences, specifically in path analysis and structural equation modeling (Skrondal & Rabe-Hesketh 2004). There, it is referred to as “Multiple indicators and Multiple Causes” (MIMIC, Jöreskog & Goldberger 1975), or as reduced-rank regression with a factor analytic structure for the residual covariance matrix (Davies & Tso 1982), but note that those developments were restricted to normally distributed responses only. Here we instead refer to the method more generally, and for non-normally distributed responses, as concurrent ordination. When there is no effect of the predictors, concurrent ordination simplifies to an unconstrained ordination, and similarly without the LV-level error and when only predictor variables are included, the model simplifies to a constrained ordination, so that it is similar to the popular constrained ordination methods CCA and RDA.

Using simulations we showed that concurrent ordination was more accurate at retrieving the latent variables and species responses than CCA, while RDA performed similarly to concurrent ordination for Gaussian responses. In CCA and RDA the latent variables are a perfect functions of the measured predictors. However, in reality it can often be unclear which predictors are important in representing the major structure of community composition, so that accounting for additional residual information is important. Indeed, accounting for residual information from species responses in an ordination explicitly addresses the concern shared by community ecologists over discarding information that the ecologist is unaware of (Palmer 1993; Økland 1996; McCune 1997; ter Braak & Šmilauer 2015).

The weighted average or weighted summation scores from CCA and RDA can be considered minimally constrained, unlike linear combination scores (Palmer 1993), but nevertheless do not formally and sufficiently account for residual information provided by species responses. This is not unexpected, as classical constrained ordination methods were designed for the purpose of filtering variation only due to the predictors. Concurrent ordination combines an unconstrained ordination with a linear regression model, so that site scores can be constructed that include a measured component due to the predictors and an unmeasured component due to the residual. Constrained ordination is often described in similar terms, but site scores are predicted excluding the error term that is an integral part of the linear regression.

It is important to note that the estimator for the canonical coefficients depends on the LV-level error in a concurrent ordination. Consequently, omitting the LV-level term, as constrained ordination does, will affect the estimates of the canonical coefficients as well as their confidence intervals. This is a similar situation to that in mixed-effects models, where omission of a random-effect in the case of (un)balanced designs and non-linear responses is known the affect the parameter estimates for fixed-effects (Searle 1971; Muff *et al.* 2016). We thus conclude that only in balanced designs (i.e., with the same number of replicates for all sites), and for normally distributed responses (i.e., as for RDA in the simulation study), the LV-level error can be omitted without inducing bias to other parameters, though even then the confidence intervals for the canonical coefficients will be too narrow. It is still unclear to us what the consequences are of omitting the LV-level error for inference on the predictor effects, and we consider formally exploring this as an avenue for future research (though see e.g., Ritz & Spiegelman 2004).

We demonstrated how to apply concurrent ordination using two example datasets, one of Swiss alpine plants (D’Amen *et al.* 2017) and another of vascular plants on the island Skabbholmen in Sweden (Cramer & Hytteborn 1987). In both examples, the presence of residual variation unaccounted for by the predictor variables demonstrated the need to account for residual variation in community ecological studies using dimension reduction techniques. We assessed importance of the predictors in the ordination using a semipartial 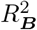 (Edwards *et al.* 2008), which can also be calculated for model-based constrained ordination if the LV-level error is excluded (though omitting the LV-level error will affect the magnitude of the 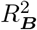 statistic). In both examples, one of the latent variables in concurrent ordination included a parameter estimate for the scale parameter of the LV-level error that was on the boundary of the feasible parameter space. For the second example the predictors were unrelated to one of the latent variables. In such situations the concurrent ordination can be simplified by omitting the LV-level error or predictors for some dimensions, and doing so is likely to improve convergence of the models (Oberpriller *et al.* 2022).

To conclude, the concurrent ordination framework provides a suitable alternative for the multivariate analysis of ecological data, with or without LV-level error. Our proposed approach provides access to standard tools for statistical inference such as statistical uncertainties of parameters estimates, model selection tools, P-values related to a Wald-statistic that can be used to determine sufficient evidence for the effect of predictors (Muff *et al.* 2022), and residual diagnostics, all of which are available as part of the `gllvm R`-package (Niku *et al.* 2020). The package also includes a vignette that demonstrate use of the proposed method through a set of worked examples. The methods presented here provide an extended version for various types of multivariate analyses, and in general have merit for the ordering of sites and species. Future research could focus on extending the method, by allowing for non-independence of the LV-level errors, e.g. due to spatial autocorrelation.

## Acknowledgements

The authors thank Cajo ter Braak and one anonymous reviewer for helpful comments on earlier drafts of the manuscript. The aspect data was provided by Olivier Brönnimann from the Spatial Ecology Group at the Univeristy of Lausanne. The species names for the Skabbholmen dataset were verified by Håkan Hytteborn and Bente Jessen Graae. Jenni Niku helped with improving the software implementation for the `gllvm R`-package. B.V. was supported by a scholarship from the Research Council of Norway (grant number 272408/F40). F.K.C.H was supported by an Australia Research Council Discovery Fellowship (grant number DE200100435).

## Authors contributions

B.V. conceived the ideas. B.V. and F.K.C.H designed the methodology. All authors contributed to the writing, reviewing and editing of the draft and gave final approval for publication.

